# Genomic divergence in allopatric Northern Cardinals of the North American warm deserts is associated with behavioral differentiation

**DOI:** 10.1101/347492

**Authors:** Kaiya L. Provost, William M. Mauck, Brian Tilston Smith

**Author notes:** Corresponding author: Kaiya L. Provost. Present address: William M. Mauck III, New York Genome Center, 101 6^th^ Ave, New York, New York 10013, USA.

## Abstract

Biogeographic barriers are thought to be important in initiating speciation through geographic isolation, but they rarely indiscriminately and completely reduce gene flow across the entire community. Understanding which species’ attributes regulate a barrier could help elucidate how speciation is initiated. Here, we investigated the association of behavioral isolation on population differentiation in Northern Cardinals (*Cardinalis cardinalis*) distributed across the Cochise Filter Barrier, a region of transitional habitat which separates the Sonoran and Chihuahuan deserts. Using genome-wide markers, we modeled demographic history by fitting the data to isolation and isolation-with-migration models. The best-fit model indicated that desert populations diverged in the mid-Pleistocene and there has been historically low, unidirectional gene flow into the Sonoran Desert. We then tested song recognition using reciprocal call-broadcast experiments to compare song recognition between deserts, controlling for song dialect changes within deserts. We found that male Northern Cardinals in both deserts were most aggressive to local songs and failed to recognize across-barrier songs. A correlation of genomic differentiation despite historic introgression and strong song discrimination is consistent with a model where speciation is initiated across a barrier and maintained by behavioral isolation.

## INTRODUCTION

Populations are frequently separated by biogeographic barriers, the ecological and physical features in the landscape that prevent organisms with particular traits from dispersing across them (Simpson, 1940; Mayr, 1963; Coyne & Orr, 2004). Geographic isolation via barriers is thought to be the dominant mode of speciation (Coyne & Orr, 2004; but see Nosil, 2008; Pinho & Hey, 2010), but not all barriers indiscriminately and completely reduce gene flow across the entire community. Filter barriers, originally conceived as filter bridges (Simpson, 1940), are a specific type of feature that preferentially allow organisms with particular traits to pass through. Using a modern example, urban environments act as a filter that preferentially allow birds with particular phenotypes to thrive (e.g., sedentary forest-dwelling birds) while excluding others (e.g., small-bodied birds with long incubation times; Croci et al 2008). Filters can be either abiotic or biotic in nature and can change over time, becoming more or less permissive (Lomolino et al., 2010). Certain populations exchange genes freely through these barriers while others show a complete cessation of gene flow, resulting in assemblages whose species have different patterns of genetic connectivity across the filter.

While the pattern of isolation by barriers is well-known, the attributes that allow organisms to pass through biogeographic filters are comparatively understudied. The meeting of taxa at distinct geographic areas of secondary contact, or suture zones, has been documented across North America and other regions (Remington, 1968; Swenson & Howard, 2004; Swenson & Howard, 2005; Swenson, 2006). For instance, avian hybrid zones are known to cluster in the Great Plains (Swenson, 2006). While suture zones have shown that gene flow can occur in some taxa (Remington, 1968), they may also provide insight into why other taxa do not show introgression during secondary contact. Possible factors include variations in dispersal ability, differences in niche breadth or preferences, pre- and post-zygotic reproductive barriers, or a combination of these. Understanding the importance of any of these mechanisms in preventing gene flow will clarify the link between the genetic structuring of populations and the subsequent initiation of speciation.

The Cochise Filter Barrier, which is a geological and ecological formation separating the Sonoran and Chihuahuan deserts in the southwestern United States and northern Mexico, is one example of the heterogeneous effects that filter barriers can have on the surrounding biota. The barrier formed during the uplift of the Sierra Madre Occidental and the Pliocene and Pleistocene glacial cycles (Morafka, 1977). Pleistocene glacial-interglacial cycles caused the Sonoran and Chihuahuan deserts to expand and contract repeatedly, alternately connecting the deserts via an arid corridor and separating the deserts with woodlands during colder glacial periods (Van Devender et al., 1984; Van Devender, 1990). Under contemporary climatic conditions, the deserts are connected by a corridor of xerophytic vegetation (Van Devender et al., 1984; Van Devender, 1990; Holmgren et al., 2007).

The genetic turnover of taxa between the Sonoran and Chihuahuan deserts generally occurs between the Baboquivari Mountains of Arizona and the Trans-Pecos of Texas (108-112 °W longitude; reviewed by Hafner & Riddle, 2011), but there is no narrow concordant transition zone across taxa (Pyron & Burbrink, 2010). Some species are genetically isolated across the Cochise Filter Barrier while others are unstructured or appear to maintain gene flow in birds (Zink et al., 2001; Zink, 2002; Riddle & Hafner, 2006) and other vertebrates (Orange et al., 1999; Riddle et al., 2000; Serb et al., 2000; Jaeger et al., 2005; Riddle & Hafner, 2006; Castoe et al., 2007; Pyron & Burbrink, 2009; Mantooth et al., 2013; Schield et al., 2015; Schield et al., 2017; Myers et al., 2017); as such, the barrier is semipermeable to gene flow. For populations distributed across the Cochise Filter Barrier, it is unclear which mechanisms have facilitated or inhibited existing gene flow. To address one such mechanism by which barriers prevent gene flow after isolation, we examined the role of behavior in a resident songbird, the Northern Cardinal (*Cardinalis cardinalis*).

In birds, a particularly salient prezygotic reproductive barrier comes in the form of song. Male songbirds sing to defend their territories and attract mates, and behave aggressively towards conspecific songs and intruders (Gill & Lanyon, 1964; Catchpole & Slater, 1995). Juveniles learn their songs from nearby singing adults (e.g., Jenkins, 1978) and thus slight variations in dialect are retained in very localized areas as a consequence of low dispersal (Lemon, 1975; Lanyon, 1979; Slater, 1989; Marler, 1997). Females can discriminate against song types and dialects to choose mates (West et al., 1981; O’Loghlen & Rothstein, 1994). Divergence in male traits may be associated with speciation, either directly through male-male competition or as mediated by female choice (Tinghitella et al., 2018; Uy et al., 2018; Burdfield-Steel & Shuker, 2018). Assessing male-male competition is more tractable than female choice in experiments on wild birds because males more readily respond to experimental stimuli. Most studies done to date assumed that males and females behave similarly to each other in terms of response to male song (e.g., Derryberry, 2011; Dingle et al., 2010). The few studies that have assessed both sexes have found no evidence that males are more discriminating than females, supporting such an assumption (Uy et al., 2018). Thus, to obtain larger samples sizes in order to tease apart the effects of ancestry and geography on song discrimination, we focused on male responses in this study.

If males on either side of a barrier sing different songs, females may not recognize a novel song as a reproductive signal, reducing interactions between populations and preventing successful gene flow (Searcy & Andersson, 1986; Hunt et al., 2009; Lipshutz, 2017b). Populations would thus become isolated and differentiation would be maintained by behavioral isolation. Like other songbirds, Northern Cardinal males are generally less aggressive in response to unfamiliar songs, and the species is sensitive to dialect changes that can occur over tens of kilometers, responding with decreased levels of aggression to more distant dialects (Lemon, 1966; Lemon, 1974; Anderson & Conner, 1985). Given this sensitivity, Northern Cardinals are likely to use song recognition as a means of species recognition, making them a candidate for investigating the relationship between genetic connectivity and song divergence. At present, however, the impact of song variation across the deserts has not been examined with respect to the impact of dialect changes due to geography. Such an assessment would disentangle the roles of dialect changes due to geographic distance vs. dialect changes due to allopatry across a barrier, sexual selection, or reproductive character displacement.

The Northern Cardinals in the Sonoran and Chihuahuan deserts are a tractable model for testing these changes as they are currently allopatric without a known contact zone. Northern Cardinals have a fragmented distribution across the Cochise Filter Barrier, being separated by a gap of ~200 km that corresponds to the elevational and environmental change of the Cochise Filter Barrier. Due to this, there should be no contemporary impact on the dialects in this region either from song learning or reinforcement. Thus, this system allows for the study of the early stages of speciation that are unbiased by later interactions in secondary contact.

Prior work has shown that this species shows phenotypic (Ridgway, 1901) and genetic (Smith et al., 2011; Smith & Klicka, 2013) differentiation across the barrier. At present, however, the amount of gene flow that occurs across the barrier, either currently or historically, is not known. There are multiple potential factors that could have led to the separation of these lineages. From a pure dispersal standpoint, Northern Cardinals should readily be able to cross this region over evolutionary time assuming no environmental or behavioral barriers. The Northern Cardinal has undergone dramatic contemporary range expansions into the northern United States and Canada and there are numerous records of individuals that have dispersed a similar distance or greater across unsuitable habitat (Halkin & Linville, 1999). Further, vagrant Northern Cardinals are regularly observed well outside their species’ resident distribution (Sullivan et al., 2009). Reconstruction of Pleistocene species distribution models also indicate that there were more suitable climatic conditions across the Cochise Filter Barrier during glacial cycles (Smith et al., 2011). Thus, it is unlikely that the observed differentiation across the Cochise Filter Barrier is due solely to limitations on this species’ dispersal capabilities. Instead, it seems more likely that sexually-selected and/or ecologically-mediated traits have impacted divergence in this species.

How focal populations respond to vocal dialects is expected to be linked to the magnitude and direction of gene flow across the Cochise Filter Barrier. Individuals who successfully migrate should exchange their genetic material and dialect with the local population. When no gene flow occurs across the barrier (i.e., pure isolation), both populations should respond aggressively to their own dialect, and should ignore the other desert’s dialect. Likewise, if gene flow occurs equally across the barrier in both directions (i.e, isolation with symmetric gene flow), then focal populations should respond equally aggressively to both their own dialect and to the other desert population’s dialect. When gene flow is biased in one direction (i.e., isolation with asymmetric or unidirectional gene flow), one population is exposed to both dialects while the other is exposed only to their own. Because of this, the population receiving more migrants should respond aggressively to both dialects. However, the population that receives fewer migrants should respond less aggressively to the foreign dialect, or even ignore it. Populations that have come into secondary contact are predicted to show equal aggression to dialects if focal populations are tested within the contact zone, and ignore foreign songs outside of the secondary contact zone.

Here, we tease apart these complex scenarios by integrating demographic modeling of genome-wide genetic variation and field-based experiments to test how a barrier regulates speciation. First, we characterized population structure and fit genomic data to pure isolation and isolation-with-migration models. From these analyses, we inferred the depth of divergence and extent of gene flow across the barrier. Second, we performed call-broadcast experiments in each desert to assess male aggression to local and non-local songs. If song discrimination is a reproductive filter, then we predict that isolation and the extent of gene flow will be correlated with male aggression to non-local songs (Figure 1). By combining genomic estimates of isolation and introgression with responses of wild birds to song differences involved in mate choice, we explore whether behavioral isolation helps regulate gene flow across filter barriers.

**Figure 1:**
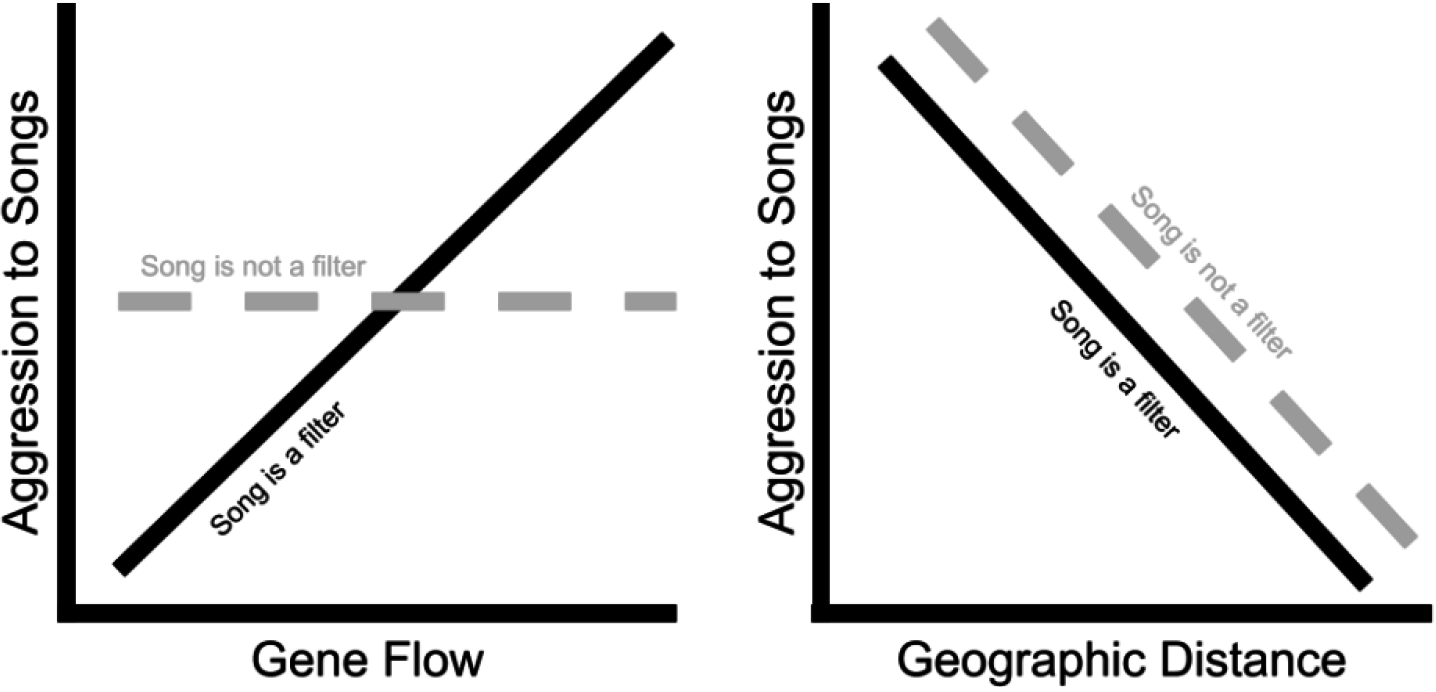
Illustration of hypothesized relationships between gene flow and song (left panel) and geographic distance and song (right panel). If song acts as a reproductive filter (solid black line), male aggression to non-local songs should be high if gene flow between their populations is high (and thus if genetic isolation is low). However, if song discrimination does not act as a reproductive filter (dashed grey line), there should be no correlation between gene flow among populations (or genetic isolation) and aggression. Irrespective of whether song acts as a filter, populations that are at larger geographic distances should show lower aggression to each others’ songs due to dialect changes.

## MATERIALS AND METHODS

### Collection of genomic data

We sequenced genome-wide markers from vouchered genetic samples collected east and west of the Cochise Filter Barrier (Figure 2). Northern Cardinals are sparse in the region of the barrier itself and as such we lack sampling there (Sullivan et al., 2009). All of the western samples occur in the Sonoran Desert (N = 54) while the eastern samples include the Chihuahuan Desert and adjacent areas in New Mexico and Texas (N = 31). For simplicity, we designated western individuals the Sonoran population and eastern individuals the Chihuahuan population, though they include individuals outside of the deserts proper. We also included three individuals of *C. cardinalis carneus* from the Pacific Coast of Mexico and one of *C. sinuatus* as outgroups. We used a Qiagen DNeasy blood and tissue kit to isolate pure genomic DNA for later sequencing following the built-in protocol with minor modifications (See Supplementary Materials for more detailed methods). Double-digest restriction-site-associated DNA sequence (ddRAD) libraries were prepared and sequenced at the University of Texas Austin Genomic Sequencing and Analysis Facility using a protocol modified from Peterson et al. (2012). The ddRAD libraries were sequenced on a single lane of an Illumina HiSeq 4000 PE 2×150, producing paired-end reads of approximately 200-300 base pairs. We processed the raw genomic data using PyRAD version 3.0.66 (Eaton, 2014; Settings: Mindepth 6, NQual 4, Wclust 0.85, Datatype pairddrad, MinCov 4, MaxSH 3, maxM 0, filter NQual+adapters, maxH 10, trim overhang 2,2). Submission of these data to the Short Read Archive and to Dryad is in progress. We characterized genetic structure with a STRUCTURE analysis (Pritchard et al., 2000; Hubisz et al., 2009), running five runs each of clusters K = 1 to K = 5 for 100,000 generations of burn-in and 500,000 generations of run time. We evaluated the best K value using both the Evanno et al. (2005) method and the Puechmaille (2016) method, as implemented in StructureSelector (Li & Liu, 2018). For the Puechmaille (2016) method, we used a mean membership threshold value of 0.5 (but see Supplementary Materials).

**Figure 2:**
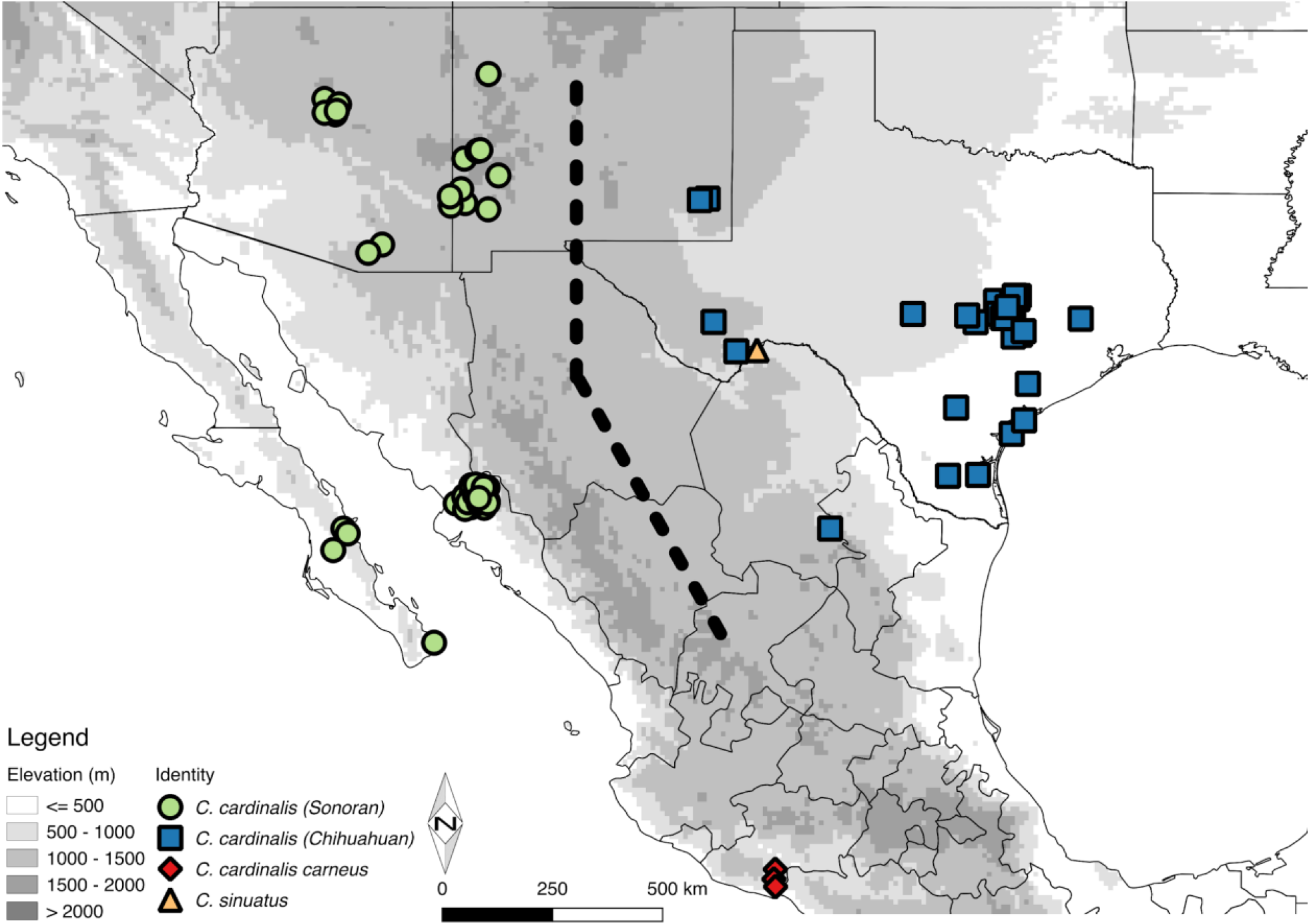
Location of vouchered *Cardinalis cardinalis* genetic samples. Points are jittered slightly to avoid overlap. Note that Chihuahuan group includes all samples east of the Cochise Filter Barrier, including individuals collected outside the Chihuahuan Desert proper. Black dotted line shows the approximate region of the Cochise Filter Barrier.

### Demographic modeling

We modeled demographic history (*N*_*E*_ = effective population size, *T* = time of divergence, and *m* = gene flow between populations) and performed model selection using fastsimcoal2 version 2.52.21 (Excoffier & Foll, 2011; Excoffier et al., 2013). Using single nucleotide polymorphism (SNP) data in variant call format (VCF) from PyRAD, we generated an unfolded joint site frequency spectrum (SFS) using ∂a∂i (Gutenkunst et al., 2009) and simulated from it in fastsimcoal2. We projected the SFS down to a smaller number of samples (10 Sonoran × 10 Chihuahuan × 3 *C. c. carneus*), averaging over missing data for each population. We then used the SFS to simulate demographic histories under multiple models, testing the fit of the simulations to our empirical SFS data, assuming a mutation rate of 2.21 × 10^−9^ mutations per site per year (*μ*, used to estimate *N*_*E*_ from *θ* = 4 *N*_*E*_ *μ*; Nam et al., 2010) and generation time of one year based on the year of first breeding of the species (Halkin & Linville, 1999). We tested six demographic models (Supplementary Figure 2) representing isolation with or without gene flow between the Sonoran and Chihuahuan deserts, where isolation refers to allopatric populations: 1) pure isolation, 2) isolation with symmetric migration, 3) isolation with asymmetric migration, 4) isolation with migration from Sonoran to Chihuahuan only, 5) isolation with migration Chihuahuan to Sonoran only, and 6) isolation with secondary contact. The models used three populations: the Sonoran group, Chihuahuan group, and the outgroup, *C. c. carneus*.

For each model, we ran 25 iterations of 100,000 simulations on 100 parameter sets, selected the iteration with the highest estimated maximum likelihood, and chose the best model by comparing small-sample-size corrected Akaike information criterion (AICc) scores. We considered a model that improved the AICc score (ΔAICc) by 2.0 to be a significant improvement, with an improvement of 10.0 or more highly significant. We then chose the best model and ran 100 bootstraps (100,000 simulations of 100 parameter sets) to calculate mean and 95% confidence interval estimates for effective population sizes, gene flow rates, time of divergence, and time of secondary contact. Note that gene flow rates are mean estimates over the entire period in which gene flow can occur, i.e., from the time of divergence (or secondary contact) to present. As such, this does not capture temporal variation in gene flow rate. Prior specifications were as follows: all effective population size estimates were a log-uniform distribution with a range between 50,000 and 1,000,000 haploid individuals, all migration estimates were a log-uniform distribution with a range between 0.001 and 20 individuals per generation, and all divergence times were a uniform distribution with a range between 100,000 and 3,000,000 generations. In addition, we constrained the time of divergence between the Sonoran and Chihuahuan deserts to be more recent than the time of divergence between the desert populations of Northern Cardinal and the *C. c. carneus* outgroup, and the time of secondary contact to be more recent than either of those values.

### Testing behavioral isolation across the Cochise Filter Barrier

We used a playback (call broadcast) experiment to examine the response of males to simulated intruders (Peters et al., 1980; Derryberry, 2007; Derryberry, 2011). These experiments assessed aggression towards a treatment, i.e., calls from one of three geographic areas and a control. Playbacks were in accordance with Columbia University’s Institutional Animal Care and Use Committee (approved as Protocol AAAM5551). We performed experiments on both Sonoran and Chihuahuan individuals in mesquite scrub habitat. Sonoran playback sites were near Portal, AZ, and Chihuahuan playback sites were in Big Bend National Park, TX. For each desert experiment, we categorized recordings into one of four treatments: Local, Distant, Across-Barrier, and Control (see Supplementary Figure 3). Local recordings came from the same population whose responses we measured (i.e., the focal population). Distant recordings came from the same desert as the focal population, but from a large geographic distance (~450–625 km). Across-Barrier recordings came from the other desert lineage, which were necessarily at a large geographic distance. Distant songs and Across-Barrier songs should have been novel to the Local population for both desert populations (Lemon, 1975). Control recordings were that of a Cactus Wren (*Campylorhynchus brunneicapillus*), which is a distantly-related bird common in both deserts. We chose these recordings to compare the effects of distance and presumed genetic relatedness on a population’s response to a recording.

**Figure 3:**
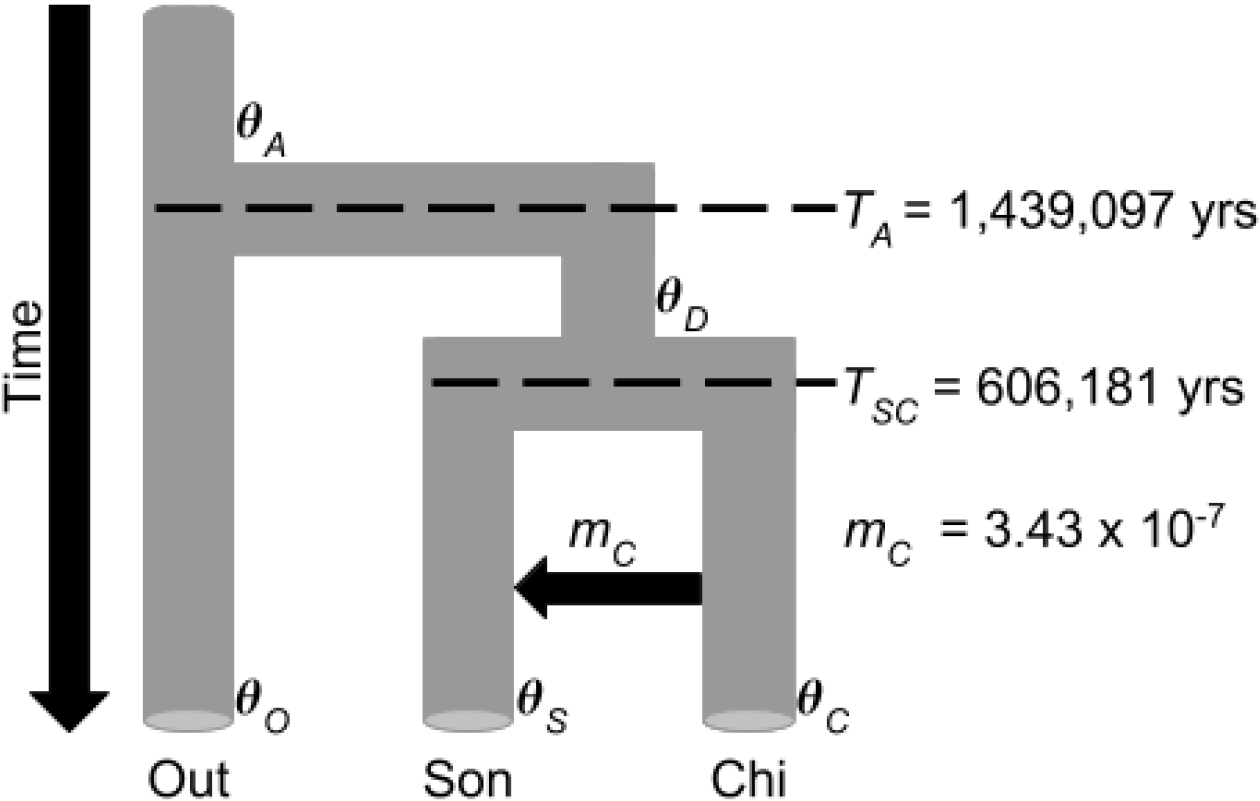
Best demographic model selected by fastsimcoal2, modified from Carstens et al. (2013); see Supplementary Figure 2 for all models tested. Population sizes given by *θ*, migration rate from Chihuahuan to Sonoran desert given by *m*. Population names Out, Son, and Chi refer to outgroup *C. c. carneus,* Sonoran Desert population, and Chihuahuan Desert population, respectively. Subscripts designate the following: *A*: *C. cardinalis* species ancestor value; *D*: Sonoran-Chihuahuan desert ancestor value; S: Sonoran value; O: outgroup *C. c. carneus* value; *C*: Chihuahuan value.

Six recordings were downloaded from Xeno-Canto (xeno-canto.org), an online open-access birdsong repository (XC141500, XC240937, XC233123, XC233122, XC211632, XC211631). All other recordings were made by the authors using a Sennheiser ME66 shotgun microphone connected to a Roland R-26 portable recorder at a sample rate of 96 kHz and a sample size of 24 bits (Xeno-Canto numbers pending). For each recording, we created a stimulus set of approximately 3 min duration (range: 2:59–3:10 min). Each stimulus set included three 50-second bouts of song, with ten-second periods of silence following each bout. We added these silent periods to mimic multiple singing bouts as Northern Cardinals in these areas rarely sang continuously for 3 min. We used 5–7 stimulus sets for each treatment except for the Control, in which we had one single Cactus Wren recording. We used an HTC6500LVW cell phone and an HMDX Jam Bluetooth speaker (model HX-P230GRA) with a ~1 m long 3.5 mm audio cable to play the stimulus sets. We placed the speaker on a tripod ~1 m off the ground. Stimulus sets were standardized to 80–85 dB as measured from 1 m away from the speaker. We broadcasted multiple stimulus sets per recording locality (or Treatment) to control for pseudoreplication (Kroodsma, 1989). The Control stimulus set was played at every site (N = 67 Sonoran, 61 Chihuahuan). We played the Sonoran stimulus sets 11–15 times each, except for two Across-Barrier stimulus sets which were played once and twice, respectively. The Chihuahuan stimulus sets were played 9–17 times each.

### Behavioral study sites

To choose sites for behavioral experiments, we placed GPS points in habitat known to have territorial males. We did not verify that points had resident males before beginning playback experiments on those sites. When males were found, we did not mark individuals but instead assumed that males found on territories were the territory holders. Sonoran Northern Cardinal territories in Pima County, AZ, average 1.56 hectares (Gould, 1961) while Chihuahuan territories in Nacogdoches County, TX, have a mean ± standard deviation of 0.64 ± 0.14 hectares (Conner et al., 1986). Territory sizes range within the species from 0.21-2.60 hectares (Halkin & Linville, 1999) with size partially dependent on foliage (Conner et al. 1986). Assuming circular territory shape, this species forms territories with radii averaging 70.5 m in the Sonoran Desert and 45.1 m in the Chihuahuan Desert (range: 25.8-91.0 m). We assumed that distances between sites were sufficient to minimize territorial overlap, and that each site comprised a single territory. Distances between Sonoran nearest-neighbor sites ranged from 38–205 m. Distances between Chihuahuan nearest-neighbors were 92–204 m. We generated transects of 6–13 sites each whose playbacks we always completed in the same order, barring dangerous conditions such as sudden inclement weather (Supplementary Table 1, Supplementary Table 2). We did this to minimize time-of-day and time-of-year effects within sites. Sites within transects were at least 90 m apart to minimize repeat testing of the same territories.

Playback experiments were done during the breeding season of 2015 for Sonoran birds and 2016 for Chihuahuan birds. We performed all playbacks at each site at nearly the same time each day, completing all trials within nine days of initiating experiments at a site. Males had at least ~24 hours between playbacks to return to a non-disturbed state. Two Chihuahuan sites form exceptions to these rules as they had multiple playbacks per day, were done substantially later in the day, and with less than an hour between playbacks.

We conducted four playbacks at each site, randomly selecting one stimulus set from each of the four Treatment localities (Local, Distant, Across-Barrier, and Control). We observed the site for 3 min before playback began (“Pre-Playback” period). We then played the 3 min stimulus set (“Playback” period), and continued to observe the male during a silent post-playback period of 3 min (“Post-Playback” period). In most analyses, we combined the Playback and Post-Playback periods *a posteriori* into a “Response” period. During each period, we recorded multiple aggression measures: 1) the number of flights greater than 1 m, or “Flybys”, within the site; 2) the presence of chip alarm calls, or “Chips”, within the site; 3) the number of male songs produced within the site, or “Close Songs”; 4) the number of male songs produced outside the site, or “Far Songs”; and 5) the distance of males from the speaker recorded every 10 seconds, or “Distance by Time”. Distance to playback equipment is a known proxy for avian aggression and mating signal recognition (Searcy et al., 2006). We categorized distances into distance bins (0–1 m, 1–2 m, 2–4 m, 4–8 m, 8–16 m, 16–24 m, and >24 m) using markers placed 8 m and 16 m from the speaker. We chose these bins as they were easily estimated by observers while also accurately capturing distance variation of male Northern Cardinals during preliminary trials. We localized birds that disappeared from sight to their nearest distance bin by sound if we could track them without ambiguity. If the localization was ambiguous, we classified the individual as >24 m (out of the site). From these distance records, we computed the minimum distance to speaker, or “Closest Distance”, for each period.

The five aggression measures were reduced to a single composite aggression measure via principal components analysis (PCA). We calculated the principal components using Response period (Playback and Post-Playback combined) data, then used the resulting loadings to calculate *a posteriori* the first principal component, PC1, for the Pre-Playback period, or “Pre-Treatment Aggression” measure. We used generalized linear mixed models (GLMMs) to determine if there was a correlation between aggression (PC1) and Treatment. We set Treatment and Pre-Treatment Aggression as fixed effects, and included the random effects of site and stimulus set in our models to control for individual site differences and differential responses to the various stimulus sets from the same locality. Each playback (unique combination of site and stimulus set) was used as a replicate. To evaluate the importance of Treatment as a predictor, we compared the outputs of: 1) a model containing all variables and 2) a model with all variables except for Treatment. We chose the model with the smallest AICc value as the best model, evaluated the significance of Treatment and Pre-Treatment Aggression predictors using a Wald χ^2^ test, and assessed model fit by calculating adjusted R^2^ values.

## RESULTS

### Raw genomic results from ddRAD pipeline

We collected 67,191 raw loci from our ddRAD pipeline. After filtering for paralogs, this was reduced to 33,626 processed loci, from which we extracted 28,798 unlinked SNPs out of a total of 361,011 variable sites and 148,370 parsimony-informative sites. Two individuals had high amounts of missing data which led to spurious assignment to groups in preliminary analyses; as such, these individuals were removed from further analyses (see Supplementary Materials). All other individuals had between 3,353–9,752 loci associated with them (mean ± standard deviation 7,000 ± 1,490, median 6,917).

### Analysis of population structure in genomic data

STRUCTURE analyses on Sonoran and Chihuahuan individuals showed strong population assignments, with Sonoran individuals always assigned to one cluster and Chihuahuan individuals never assigned to that cluster (Figure 4). The highest log-likelihood support values were for K = 2 (mean ± standard deviation −181,543 ± 56) and K = 3 (−186,576 ± 131) populations. Comparisons of ΔK (Evanno et al., 2005) show highest support for K = 3. However when K = 3, the third cluster never achieves more than 25% assignment probability in any individual. Using the methodology suggested by Puechmaille (2016) implemented in Structure Selector (Li & Liu, 2018), the highest support for all metrics is K = 2.

**Figure 4:**
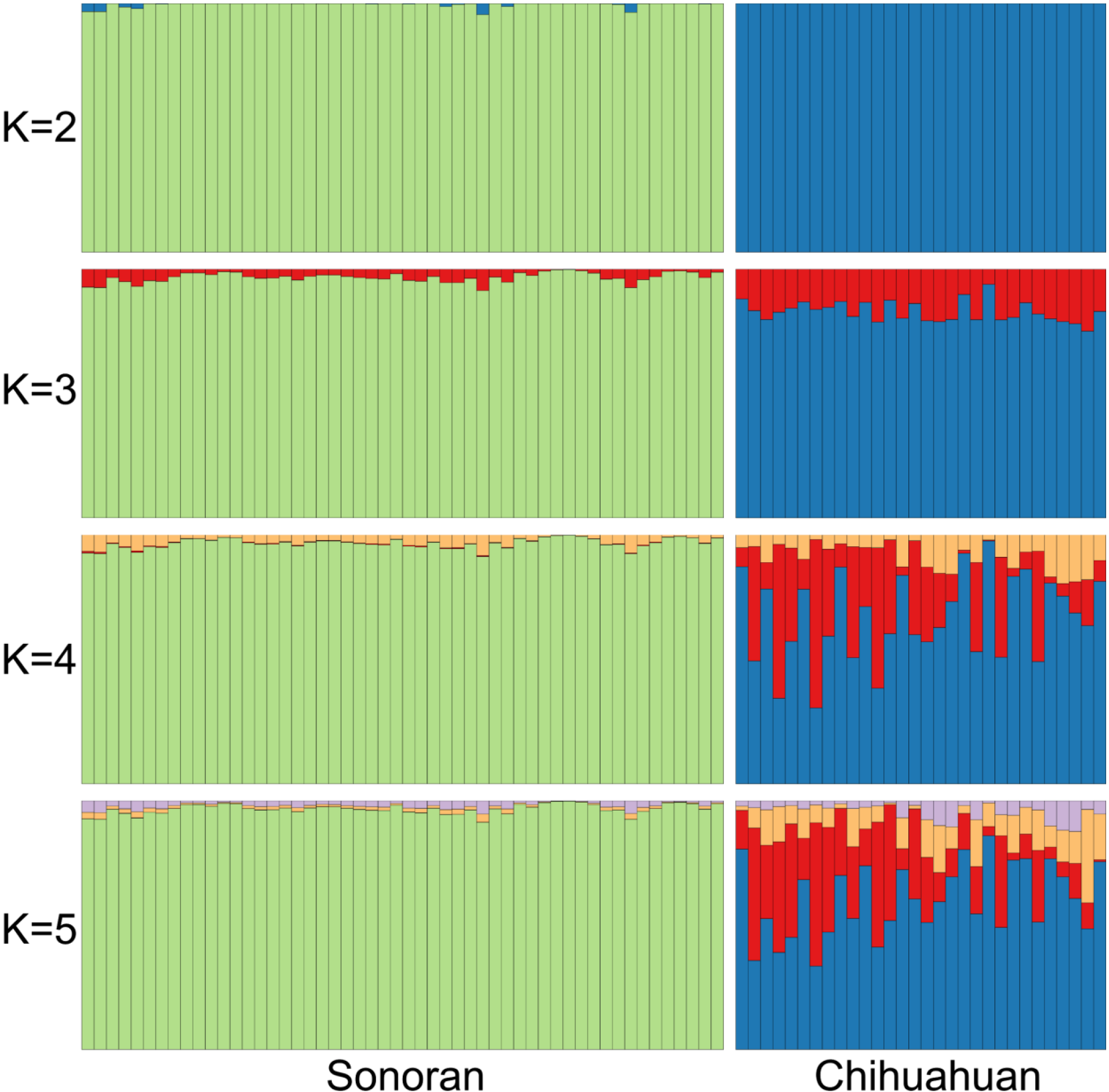
Individuals east and west of the Cochise Filter Barrier are consistently assigned to different populations. Results from STRUCTURE runs K = 2-5 are presented. Each vertical bar represents an individual bird, with the proportion of each color indicating assignment to that population.

We found that isolation with unidirectional gene flow (Figure 3) from the Chihuahuan Desert to the Sonoran Desert was the best fit model. All models with ΔAICc scores of 10.0 or less were from this demographic scenario (best model ΔAICc scores range 0.0–16.1, other model ΔAICc scores range 25.4–114.3) After bootstrapping the best model, we dated the divergence between the Sonoran and Chihuahuan populations to be a mean of 606,181 years (95% CI 583,982–600,982). The divergence between the Sonoran-Chihuahuan ancestor and the *C. c. carneus* lineage was dated at a mean of 1,439,097 years (95% CI 1,379,287–1,436,750). The Sonoran population effective population size (mean 169,621; 95% CI 165,910–170,650) was substantially smaller than the Chihuahuan (mean 839,453; 95% CI 794,972–815,171). Despite a model with gene flow receiving the highest support, the actual estimated gene flow rate into the Sonoran Desert was minute (mean 3.43 × 10^−7^; 95% CI 3.19 × 10^−7^−3.31 × 10^−7^). The support for a model with gene flow over a model of secondary contact suggests that gene flow is not recent, and more likely to reflect the presence of historical introgression.

### Levels of aggression to playbacks from different localities

We ran playback experiments at 67 and 61 sites in the Sonoran and Chihuahuan deserts, respectively, resulting in 512 total playback trials (four playbacks per site). Of these, we did not detect males at 16 Sonoran and 25 Chihuahuan sites. These sites were removed, leaving 51 Sonoran and 36 Chihuahuan sites, for 348 total playbacks. When we performed our PCA on the aggression data, the first principal component (PC1, aggression) explained 58.2% of the variation in the Response period data for the Sonoran individuals and explained 52.4% of variation in the Response period for the Chihuahuan individuals.

We compared the effects of GLMMs with and without recording locality (Treatment) as a predictor and found that Treatment had a significant effect on male aggression in both deserts (Table 1). For playbacks on Sonoran individuals, the model with Treatment was a better fit to the aggression data than the model without (ΔAICc = 9.97), though both models had equivalent adjusted R^2^ values (0.90). Wald tests indicated that both Treatment and Pre-Playback Aggression were significant factors in the full model (Treatment p-value < 0.001; Pre-Treatment Aggression p-value < 0.001). Sonoran males were significantly more aggressive to Local stimulus sets than to those from any other location (all p-values < 0.001; Figure 5). By contrast, there were no significant differences in aggression across Distant, Across-Barrier, or Control stimulus sets (all p-values ≥ 0.35).

**Table 1:** Models including Treatment locality explain responses to song dialects better than models without. Model parameter results from generalized linear mixed models for both Sonoran and Chihuahuan experiments. Treatment localities are as follows: Con = Control, Acr = Across-Barrier, Dis = Distant, Loc = Local. Pre-Agg = Pre-Playback Aggression. S.E. = Standard error.

|  | Sonoran Experiment |  | Chihuahuan Experiment |  |
| --- | --- | --- | --- | --- |
|  | Null Model† | Full Model‡ | Null Model† | Full Model‡ |
| **AICc (ΔAICc)** | 588.7 (9.96) | 579.01 (0.00) | 352.87 (4.79) | 348.08 (0.00) |
| **adjR<sup>2</sup> (R<sup>2</sup>)** | 0.90 (0.90) | 0.90 (0.90) | 0.70 (0.70) | 0.66 (0.67) |
| **Site Variance** | 0.04 | 0.04 | 0.00 | 0.00 |
| **Stimulus Set Variance** | 0.14 | 0.03 | 0.40 | 0.07 |
| **Residual Variance** | 0.34 | 0.33 | 1.15 | 1.26 |
| **Intercept $\alpha \pm$ S.E.** | 1.02 ± 0.13 | 0.60 ± 0.21 | -0.56 ± 0.20 | -0.61 ± 0.33 |
| **Intercept T (p-val)** | 7.69 (1.47 x 10<sup>-14</sup>) | 2.79 (0.0054) | 2.84 (0.0045) | -1.84 (0.065) |
| **Pre-Agg $\beta \pm$ S.E.** | 0.39 ± 0.08 | 0.39 ± 0.08 | -0.70 ± 0.12 | -0.71 ± 0.12 |
| **Pre-Agg T (p-val)** | 4.93 (8.15 x 10<sup>-7</sup>) | 5.06 (4.36 x 10<sup>-7</sup>) | 5.88 (4.05 x 10<sup>-9</sup>) | -5.81 (6.40 x 10<sup>-9</sup>) |
| **Con vs Acr $\beta \pm$ S.E.** | N/A | 0.10 ± 0.23 | N/A | 0.89 ± 0.40 |
| **Con vs Acr T (p-val)** | N/A | 0.45 (p = 0.65) | N/A | 2.22 (0.027) |
| **Con vs Dis $\beta \pm$ S.E.** | N/A | 0.18 ± 0.24 | N/A | 1.57 ± 0.40 |
| **Con vs Dis T (p-val)** | N/A | 0.78 (p = 0.44) | N/A | 3.92 (9.00 x 10<sup>-5</sup>) |
| **Con vs Loc $\beta \pm$ S.E.** | N/A | 1.09 ± 0.24 | N/A | 1.60 ± 0.40 |
| **Con vs Loc T (p-val)** | N/A | 4.65 (p = 3.27 x 10<sup>-6</sup>) | N/A | 4.00 (6.14 x 10<sup>-5</sup>) |
| **Acr vs Dis $\beta \pm$ S.E.** | N/A | 0.078 ± 0.16 | N/A | 0.67 ± 0.32 |
| **Acr vs Dis T (p-val)** | N/A | 0.48 (p = 0.63) | N/A | 2.10 (0.036) |
| **Acr vs Local $\beta \pm$ S.E.** | N/A | 0.99 ± 0.16 | N/A | 0.71 ± 0.32 |
| **Acr vs Loc T (p-val)** | N/A | 6.07 (p = 1.26 x 10<sup>-9</sup>) | N/A | 2.21 (0.027) |
| **Dis vs Loc $\beta \pm$ S.E.** | N/A | 0.91 ± 0.16 | N/A | 0.04 ± 0.32 |
| **Dis vs Loc T (p-val)** | N/A | 5.53 (p = 3.13 x 10<sup>-8</sup>) | N/A | 0.12 (0.91) |

†Null model formula: Aggression ~ Pre-Playback Aggression + 1|Stimulus Set + 1|Site. Pre-Playback Aggression is a fixed effect, while 1|Stimulus Set and 1|Site are random effects.
‡Full model formula: Aggression ~ Treatment + Pre-Playback Aggression + 1|Stimulus Set + 1|Site. Pre-Playback Aggression and Treatment are fixed effects, while 1|Stimulus Set and 1|Site are random effects.

**Figure 5:**
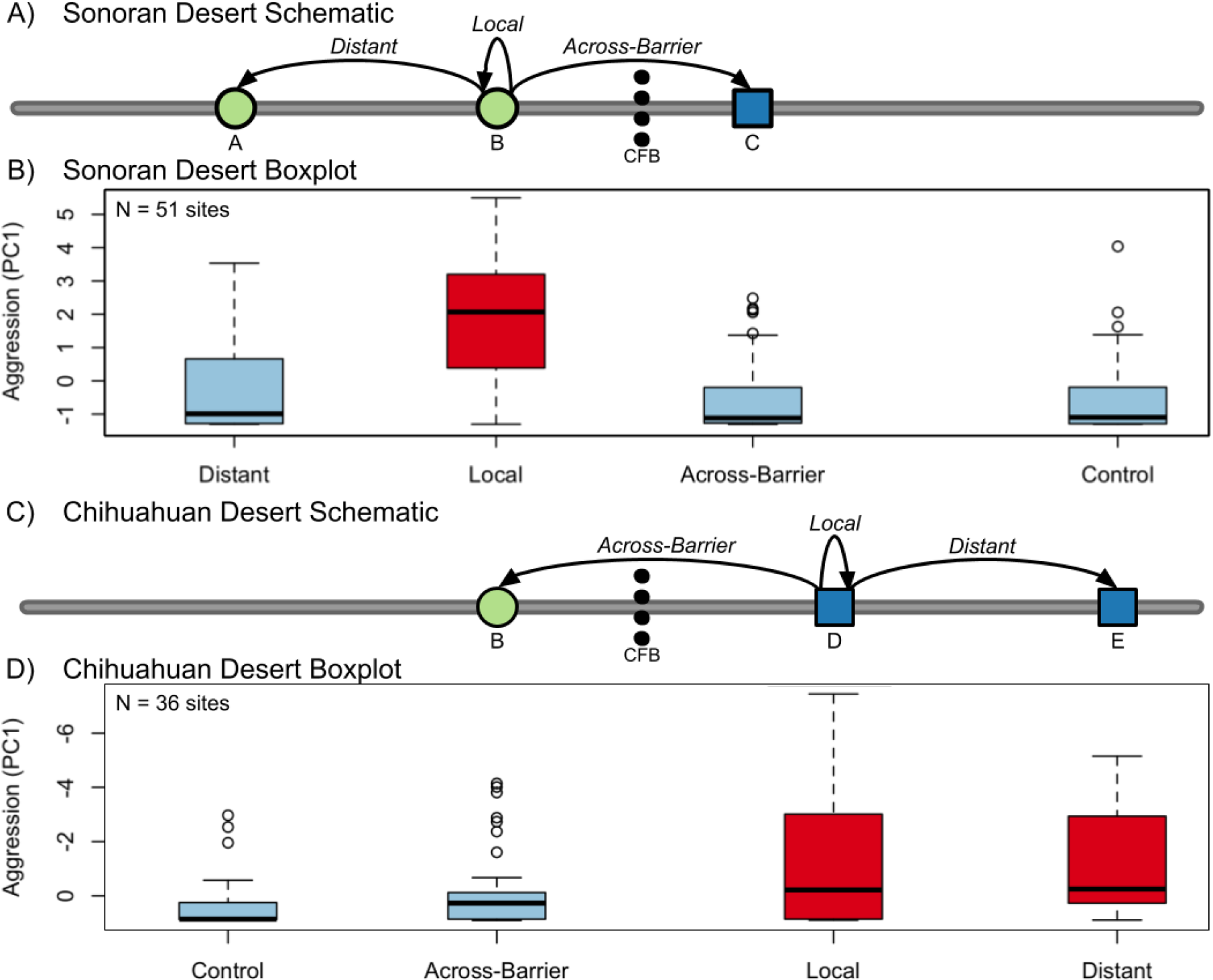
Male Northern Cardinals in both deserts were most aggressive to local songs. Panels combine schematic of playback experiment (A, C) with boxplots showing mean aggression scores (B, D). Sonoran Desert results shown in Panels A and B. Chihuahuan Desert results show in Panels C and D. Panels A and C: circles show localities within the Sonoran Desert population, squares show localities within the Chihuahuan Desert population. Distances between localities represent longitudinal distance. Arrows point from focal population of each experiment to location of Local, Distant, and Across-Barrier recording treatments (Control treatment is omitted). Letters indicate locality names: A) Western Arizona: Bill Williams River National Wildlife Refuge, AZ, Santa Maria River National Wildlife Refuge, AZ, and Wenden, AZ; B) Portal, AZ; C) Rattlesnake Springs Reserve, NM; D) Big Bend National Park, TX; E) Eastern Texas: Balcones Canyonlands National Wildlife Refuge, TX, and Austin, TX. Panels B and D: Boxplots show mean aggression scores (PC1, y-axis) for each song treatment (x-axis). Values at the top of the graph indicate higher aggression. Sites without detections were excluded. Red bars are statistically significantly different from blue bars. Note that because Chihuahuan PC1 is negatively correlated with aggression, Chihuahuan y-axis is inverted with more negative PC1 values toward top of graph (Sonoran PC1 is positively correlated with aggression).

For playbacks on Chihuahuan individuals (Table 1), AICc scores supported the model with Treatment over the model without (ΔAICc = 4.79), though the model without Treatment had greater explanatory power according to adjusted R^2^ (0.71 vs. 0.67). Nevertheless, Wald tests indicated that both Treatment and Pre-Treatment Aggression were significant factors in the full model (Treatment p-value = 0.027; Pre-Treatment Aggression p-value < 0.001). Chihuahuan males were significantly more aggressive to Local and Distant stimulus sets than to Across-Barrier or Control stimulus (all p-values ≤ 0.05; Figure 5). Local and Distant were not statistically different, and neither were Across-Barrier and Control (all p-values ≥ 0.5).

Both Sonoran and Chihuahuan birds shifted their behavioral response across the experimental periods (i.e., Pre-Playback, Playback, and Post-Playback) (Figure 6). Sonoran individuals hearing the Local treatment, and Chihuahuan individuals hearing either the Local or Distant treatment, were substantially closer to the speaker during the Playback and Post-Playback periods compared to the Pre-Playback periods. There were no significant differences between Pre-Treatment Aggression values for Treatment, Population Tested, or their interaction (all p-values ≥ 0.17). In contrast, all values were highly significant when analyzing PC1 values (all p-values < 0.001). The significant interaction indicates that there were significant differences between the deserts in the response to a treatment. Sonoran individuals are significantly more likely to be aggressive to Local songs than Chihuahuan individuals (p < 0.001), though there are not significant differences for the other three treatments (all p-values ≥ 0.17). Across all of these analyses, male Northern Cardinals in both deserts show minimal aggression toward Across-Barrier songs, treating the songs of a foreign desert as heterospecific.

**Figure 6:**
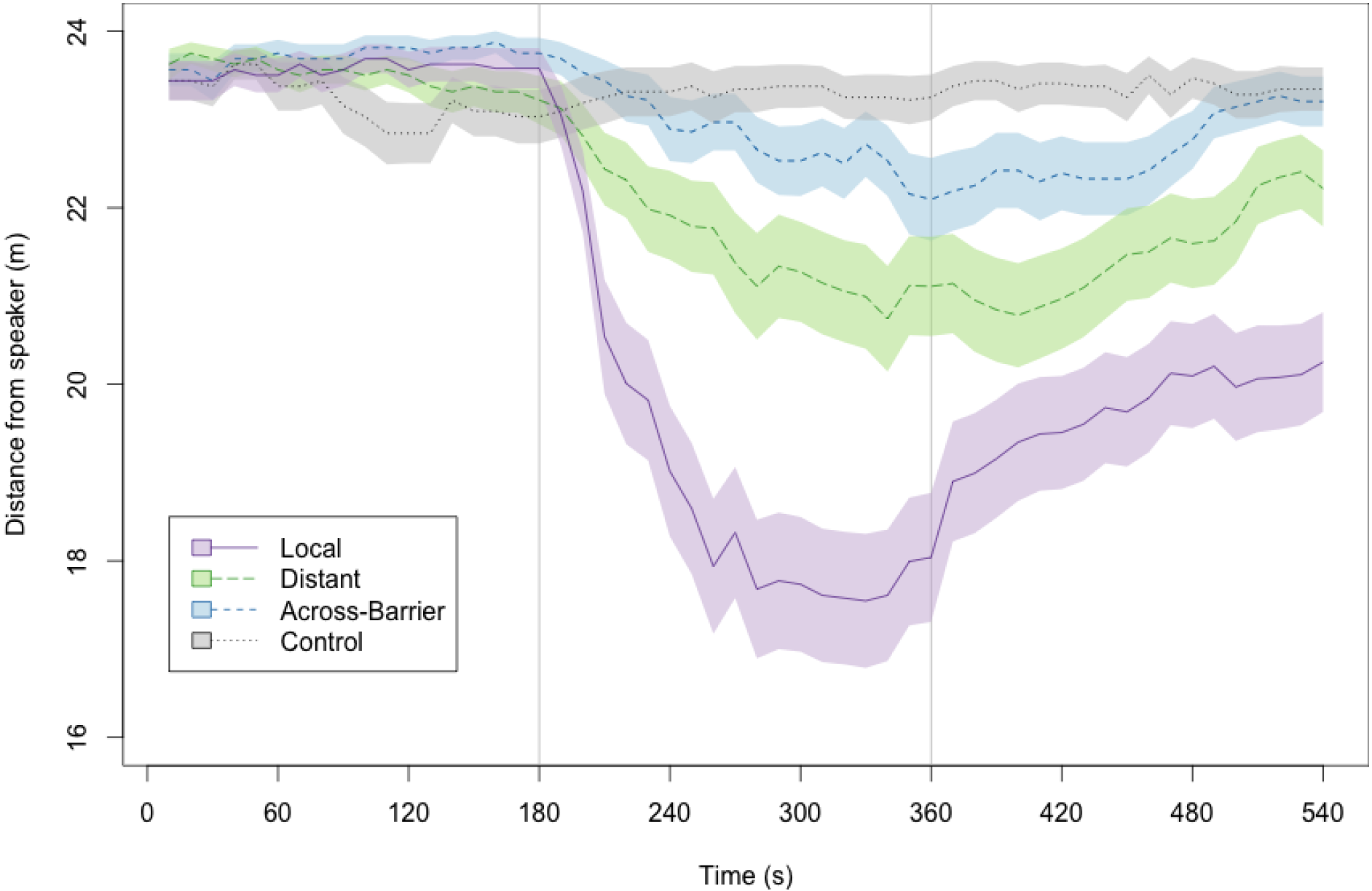
Individuals increase aggression when playbacks begin. Mean distance from speaker (m) over time for each treatment, with smaller distance values indicating more aggression. Data from both deserts were combined (N = 87 sites). Lines indicate mean distance values across all treatments, with shading around lines indicating one standard error around points. Solid purple line indicates Local treatment, large-dashed green line indicates Distant treatment, medium-dashed blue line indicates Across-Barrier treatment, and dotted grey line indicates Control treatment. Vertical lines separate Pre-Playback (left), Playback (middle), and Post-Playback (right) periods.

## DISCUSSION

Speciation across barriers and the progression from allopatric populations to genomic isolation have been a well-explored topic (Price, 2008). However, the processes that maintain low gene flow between isolated populations still prove a challenging problem in biology for understanding the early stages of speciation. Behavioral reproductive isolating mechanisms, like song differences, are expected to be important for reducing gene flow between populations, particularly when accompanied by ecological or morphological changes (van Doorn et al., 2004; Kraaijeveld et al., 2011; Lamichhaney et al., 2017). Here we found that the Northern Cardinal showed low gene flow and strong male song discrimination across the Cochise Filter Barrier. Further, we showed that song discrimination between-deserts is greater than song discrimination within-deserts, indicating that these birds show divergence in song beyond what would be expected through dialect changes alone. This strong discrimination against songs from an allopatric population may mediate the degree of gene flow permeability among deserts. Analogous studies to this one frequently examine differences between groups that are adjacent to one another (e.g., Dingle et al., 2010; Lipshutz et al., 2017), allowing for tests along axes of sympatric vs. parapatric or within vs. outside a hybrid zone. However, Northern Cardinals have no known (contemporary) contact zone across the Cochise Filter Barrier. The two allopatric populations have high connectivity within deserts and no or low connectivity between deserts, though the latter could still have been historically high in short bursts. This is in contrast to a hybrid zone system in which connectivity is relatively high in the localized area of contact. Instead of focusing on what occurs during secondary contact, our two allopatric populations allow us to explore what occurs during periods of geographic isolation. Our approach thus controls for dialect changes over large distances without relying on contact zones as proxies for connectivity.

### Potential modes of speciation across the Cochise Filter Barrier

We found that song discrimination and gene flow between allopatric populations were correlated. One potential relationship between these two factors is that song could act as a driver of divergence, with or without allopatry (Uy et al., 2018). The song differences observed in this study may have directly caused genomic divergence through assortative mating during periods of contact (Coyne & Orr, 2004; Price, 2008; Bensch et al., 1998), or perhaps reinforced existing divergence (Lynch & Baker, 1994; Grant & Grant, 1996; Lachlan & Servedio, 2004; Hoskin et al., 2005; Mason et al., 2017). Relative to the accumulation of novel genetic markers or other traits, behavioral evolution via exchange of learned song can be rapid (West-Eberhard, 1983; Allender et al., 2003; Duckworth, 2009). Further, species that learn their song evolve new dialects particularly quickly (Mason et al., 2017; but see Freeman et al., 2017). Rapid evolution of songs could mediate and/or supplement divergences across the Cochise Filter Barrier.

While we found evidence for a correlation between behavioral and genomic differentiation in this system, this does not rule out other mechanisms contributing to the divergence between the Sonoran and Chihuahuan populations. Ecological factors could have operated in tandem with behavioral factors in generating the divergence patterns seen. We dated the divergence between the Sonoran Desert and Chihuahuan Desert Northern Cardinals at approximately 600,000 years. For comparison, estimates for bird species ages in the region typically date to the Pleistocene (Smith et al., 2017). Divergence dates across the Cochise Filter Barrier, for various taxa, range from 500,000-5,000,000 years (Bryson et al., 2011; Bryson et al., 2012; Klicka et al., 2016; Zink & Blackwell-Rago, 2000; Wilson & Pitts, 2010; Smith & Klicka, 2013; Weyandt & Van Den Bussche, 2007; Leache & Mulcahy, 2007; O’Connell et al., 2017; Pyron & Burbrink, 2009; Myers et al. 2017), suggesting that population connectivity and isolation has been dynamic in the region. For Northern Cardinals, ecological niche models suggest that range shifts may have occurred during glacial cycling as the desert habitats expanded and contracted during the Pleistocene (Smith et al., 2011; Lang & Wolff, 2011; Mudelsee & Schulz, 1997). The unidirectional nature of the gene flow from the Chihuahuan into the Sonoran may also be explained by this pattern, as these niche models reveal that the Sonoran Desert was suitable habitat for birds occurring east of the Cochise Filter Barrier during historical and contemporary periods, but not the reverse, potentially due to climate and habitat differences (Smith et al., 2011; Shreve, 1942).

Demographic models show significantly higher support for a model with gene flow over a model of pure isolation, but the amount of gene flow was low. Our finding of minute mean gene flow between these populations across the years since divergence is agnostic to the timing of gene flow itself, whether it was low and protracted or high and abbreviated. The failure to support secondary contact models may also suggest that the gene flow happened earlier, rather than later, in the populations’ histories, and rejects the notion that contemporary introgression is occurring. Overall, the low amount of introgression found suggests that while gene flow was possible across deserts, it was limited either in duration or in magnitude. This suggests that some isolating mechanism evolved between the populations diverging and completely cutting off gene flow before contemporary periods. Given our findings of rapid evolution of song discrimination within the Sonoran Desert (see below), it is likely that prezgotic isolation evolved early in the differentiation of the two desert groups.

Male Northern Cardinals, irrespective of desert, do not recognize Across-Barrier dialects as conspecific. Under our tested hypothesis, this implies a pure isolation model of gene flow. Contrary to this, however, we found the presence of minute unidirectional gene flow. This result is sensible if contemporary gene flow is non-existent, as we assert above, and if song dialects evolve rapidly (e.g., West-Eberhard, 1983; but see Noad et al., 2000 for song exchange without gene flow). The observation that Sonoran birds also do not recognize Distant songs as conspecific does not contradict this finding, though all explanations as to why this population shows such behavior are speculative. These findings may be due to differential gene flow between subpopulations (e.g., McDonald et al., 2001; Rosenfield & Kodric-Brown, 2003; Grava et al., 2012; Robbins et al., 2014; While et al., 2015; Lipshutz, 2017a).

The ability of the Sonoran population to discriminate from other Distant Sonoran songs reinforces the view that prezygotic isolation can evolve before genetic differentiation. The overall strength of song discrimination of Across-Barrier dialects is consistent with reproductive isolation, and this type of finding is often used to delimit species based on biological species concepts (e.g., Caro et al. 2013; Cadena et al. 2016). However, we recommend caution, as even though it is defensible to assume that male response and female choice are tightly coupled, we did not test female choice. The few experiments done testing female choice in Northern Cardinals have not examined whether they discriminate against different genetic lineages (e.g., Yamaguchi, 1999), and so understanding their responses forms one of the critical next steps for this system. Anecdotally, during our experimental trials female Northern Cardinals occasionally responded to playbacks (N = 9 trials). These females typically behaved similarly to the focal male of the trial, and appeared more likely to investigate the speaker when hearing the local dialect, rather than a novel dialect, tentatively suggesting that males and females behave similarly in this regard (author’s unpublished data).

All in all, the Cochise Filter Barrier structures Northern Cardinal populations both genetically and phenotypically, in particular with regard to song. Given our findings, the barrier appears to be facilitated, at least in part, by strong dialect differences that have evolved between the Sonoran and Chihuahuan deserts. These dialect differences affect song discrimination in male Northern Cardinals more potently than would be expected from geographic distance alone. Across the entire bird community, it is likely that different species have developed greater or fewer dialect differences across the barrier, which may be impacting the observed genetic semipermeability. We suggest that the song discrimination and genetic divergence we found in this species directly interact with each other to create the pattern of differentiation we see across the Cochise Filter Barrier. Future studies of birds codistributed across this barrier may find similar evidence for this pattern, and investigating many different mechanisms across multiple species at once would give insight into how the Cochise Filter Barrier, and other such barriers, worked to create the species diversity seen today.

## Contributions

K.L.P. and B.T.S. conceived the study. K.L.P. and W.M.M. performed lab work. K.L.P. and B.T.S. performed behavioral work. K.L.P. ran analyses with input from other authors. K.L.P. drafted the manuscript. All authors edited the manuscript.

## Acknowledgements

We are grateful to G. Voelker (Texas A&M), S. Birks, J. Klicka (Burke Museum), P. Sweet, T. Trombone (AMNH), C. Witt, A. Johnson, and M. Andersen (Museum of Southwestern Biology) for providing or loaning the tissues used in this study. We thank the Southwestern Research Station and Big Bend National Park. This work was funded by E3B at Columbia University, the Frank M. Chapman Memorial Fund, the American Ornithological Society, the Society for Systematic Biologists, and the Linda J. Gormezano Memorial Fund. We thank M. Cords and J. Cracraft for reviews. We also thank D. Fletcher, L. Musher, L. R. Moreira, J. Koch, G. Seeholzer, E. Myers, V. Ramesh, J. T. Merwin, G. Rosen, L. Garetano, and J. McKay for helpful contributions. Lastly, we thank C. Raxworthy, A. DeRenzis, J. Arenson, A. Arteaga, P. McKenzie, and L. Bryan.

## Data accessibility

Songs used to create treatments are in the process of being uploaded to Xeno-Canto. Sequences from ddRAD procedure will be uploaded to the NCBI Short Read Archive. All data and scripts used to perform genetic and behavioral analyses will be uploaded to Dryad.

